# Rhythmic lipid and gene expression responses to chilling in panicoid grasses

**DOI:** 10.1101/2023.09.29.560160

**Authors:** Sunil K. Kenchanmane Raju, Yang Zhang, Samira Mahboub, Daniel W. Ngu, Yumou Qiu, Frank G. Harmon, James C. Schnable, Rebecca L. Roston

## Abstract

Chilling stress threatens plant growth and development, particularly affecting membrane fluidity and cellular integrity. Understanding plant membrane responses to chilling stress is important for unraveling the molecular mechanisms of stress tolerance. Whereas core transcriptional responses to chilling stress and stress tolerance are conserved across species, the associated changes in membrane lipids appear to be less conserved, as which lipids are affected by chilling stress varies by species. Here, we investigated changes in gene expression and membrane lipids in response to chilling stress during one diurnal cycle in sorghum (*Sorghum bicolor*), Urochloa (browntop signal grass, *Urochloa fusca*) (lipids only), and foxtail millet (*Setaria italica*), leveraging their evolutionary relatedness and differing levels of chilling-stress tolerance. We show that most chilling-induced lipid changes are conserved across the three species, while we observed distinct, time-specific responses in chilling-tolerant foxtail millet, indicating the presence of a finely orchestrated adaptive mechanism. We detected diurnal rhythmicity in lipid responses to chilling stress in the three grasses, which were also present in Arabidopsis (*Arabidopsis thaliana*), suggesting the conservation of rhythmic patterns across species and highlighting the importance of accounting for diurnal effects. When integrating lipid datasets with gene expression profiles, we identified potential candidate genes that showed corresponding transcriptional changes in response to chilling stress, providing insights into the differences in regulatory mechanisms between chilling-sensitive sorghum and chilling-tolerant foxtail millet.

**Significance Statement:** Plants respond to low-temperature stress in myriad ways. While core transcriptional changes are conserved across species, specific adaptive strategies do exist. However, membrane lipid responses during chilling do not appear to be conserved. Here, we collected samples from control and chilling stress–treated seedlings [PSC4] to assess gene expression and membrane lipids in three panicoid grasses to show that lipid metabolic changes follow a daily rhythm. Lipid changes in chilling-tolerant foxtail millet occurred at specific time points, partly explaining the difficulty in finding conserved chilling-induced lipid changes in previous reports. We identified specific orthologs in sorghum and foxtail millet that showed a correlation between gene expression and lipid metabolic changes; these orthologs may be used as potential target genes for developing chilling-tolerant sorghum varieties.

## INTRODUCTION

Climate change has increased the frequency and severity of extreme weather events, threatening future food supply (1). Major crop species important for global food security, such as maize (*Zea mays*), sorghum (*Sorghum bicolor*), and rice (Oryza sativa), are sensitive to chilling stress owing to their tropical origin, limiting their geographical distribution and productivity in temperate climates (2, 3). Chilling stress is major stress experienced by plants during their lifecycle, hindering energy metabolism and growth, most notably by reducing the activity of enzymes associated with photosynthesis and the energy-demanding production of protective proteins and substances (4, 5). Moreover, plants must endure daily and seasonal temperature fluctuations and unexpected extreme variations (6). Plants have devised various strategies to cope with such environmental challenges.

At the cellular level, low-temperature stress leads to increased membrane rigidity and impaired containment of cytosolic contents, resulting in cell death (7, 8). Changes in glycerolipids, major components of cell membranes, include membrane lipid polyunsaturation (9, 10), changing the ratio of lipid head groups, and removing membrane-destabilizing lipids in response to low temperature (11, 12). The contribution of unsaturated fatty acids to membrane fluidity at different temperatures and their role in protecting the photosynthetic machinery from photoinhibition under chilling stress are well known (13). However, across species, no consistent changes in membrane lipid abundance during chilling stress have been reported. These discrepancies in changes in lipid compositions or content during stress may be due to differences in the duration and/or intensity of the applied stress, time-of-day effets, and/or genetic and physiological differences across species (14).

Plant responses to these stressful environments can vary greatly at the transcriptional level, although a core set of transcriptional responses is mostly conserved across species (14). Notably, most studies of cold tolerance in the Pooideae grass subfamilyof the Poaceae (including wheat [*Triticum aestivum*], barley [*Hordeum vulgare*], and rye *[Secale cereale*]) have revealed chilling adaptive mechanisms that are not shared by closely allied subfamilies within the grasses, such as the Ehrhartoideae (which includes rice). This lack of conservation suggests that different plant lineages have adapted to growth in temperate environments using distinct genetic and physiological mechanisms. Panicoid grasses, comprising many important crops such as maize, sugarcane (Saccharum officinarum), switchgrass (*Panicum virgatum*), sorghum, and foxtail millet (*Setaria italica*), exhibit a range of sensitivities to cold temperatures (14–16). The repeated acquisition and loss of chilling tolerance within this subfamily (17, 18) make it an ideal system to study the conserved and species-specific adaptation strategies for chilling tolerance.

Sorghum, an important crop in the arid and semi-arid region of the world, originated in the semi-arid tropics of Africa and quickly spread into other parts of the world, including India, China, and the United States (19). Due to its tropical origin, sorghum is susceptible to chilling (20). While landraces and wild relatives are important gene pools for adaptive traits such as biotic stress resistance and abiotic stress tolerance (21), the limited availability of standing genetic variation and newer cropping environments require the transfer of stress adaptation mechanisms from closely related stress-adapted species. Like sorghum, foxtail millet is also a grain crop domesticated from a panicoid grass. However, foxtail millet was initially domesticated in northeast China from a wild grass, green foxtail (*Setaria viridis*) that grows in temperate climates where low-temperature stress is more common (22–24).

Orthologous genes, even within closely related taxa, can show differential regulation of chilling stress responsive gene expression between maize and sorghum, or between maize, sorghum and eastern gamagrass (*Tripsacum dactyloides*) (25, 26), suggesting that orthology alone is not a reliable predictor of stress-induced gene expression in related species (27). It can therefore be challenging to narrow down target genes for chilling-stress tolerance in sorghum and related chilling-sensitive species.

To overcome these challenges, we designed a time-course experiment to account for potential time-of-day variation and tested the relationship between chilling-stress tolerance, changes in membrane glycerolipid contents, and evolutionary relatedness using three panicoid grasses. Browntop signal grass (*Urochloa fusca*, Urochloa hereafter) is a grass closely related to foxtail millet that is less chilling tolerant. Urochloa and sorghum are more distantly related and have similar susceptibility to chilling stress. In this study, we profiled the changes in membrane lipid contents and composition and in transcript levels in these three species using paired time-course measurements of control and chilling-stress conditions and identified genes whose expression is correlated with changes in lipid composition during chilling-stress. We show that integrating lipid abundance with gene expression profiles in chilling-sensitive sorghum and chilling-tolerant foxtail millet allowed the identification of genes with previously known effects on chilling stress tolerance and novel genes with correlated expression patterns with chiling-induced changes in lipid content and composition. These genes have potential application in engineering chilling tolerance in sorghum and related chilling-sensitive grasses.

## RESULTS

### Foxtail millet is chilling tolerant compared to other panicoid grasses

Chilling stress causes phase transitions in biological membranes of cold-susceptible plants. These changes in membranes cause abnormalities in respiration and photosynthetic CO_2_ and O_2_ exchange rates (2, 28). Lower photosynthesis for prolonged periods of time, continuing for hours or days, is an essential identifier of chilling susceptibility (28). Here, we used, CO_2_ assimilation rates to quantitatively assess differences in chilling tolerance among closely related panicoid grasses (**Figure 1A**) (25). Accordingly, we took measurements on 12-day-old seedlings grown under control conditions (29°C during the day and 23°C at night) and after exposure to chilling treatment (6°C) in growth chambers for one or eight days. After eight days of chilling stress, sorghum, Urochloa, and maize showed lower values for CO_2_ assimilation, compared to the control time point, indicating impaired photosynthetic activity. In fact, sorghum and Urochloa seedlings had dead leaves which was reflected in the negative CO_2_ assimilation values (**Figure 1B**). Foxtail millet showed moderate impairment in its photosynthetic rate as its CO_2_ assimilation measurements remained at about 55% of control levels even after eight days of stress, indicating higher tolerance to chilling (**Figure 1B**) consistent with its native range and center of domestication in Northern China (23, 29). Based on these photosynthetic measurements, we classified the four panicoid species into two categories: chilling-susceptible - sorghum, Urochloa, and maize, and chilling-tolerant - foxtail millet. Prolonged stress clearly differentiated tolerance levels in foxtail millet. Urochloa and sorghum seedlings were dead after two days of recovery at 30°C, following two weeks of chilling stress at 6°C, while foxtail millet seedlings looked healthier with fewer necrotic leaves (**Figure 1C**).

**Figure 1.**
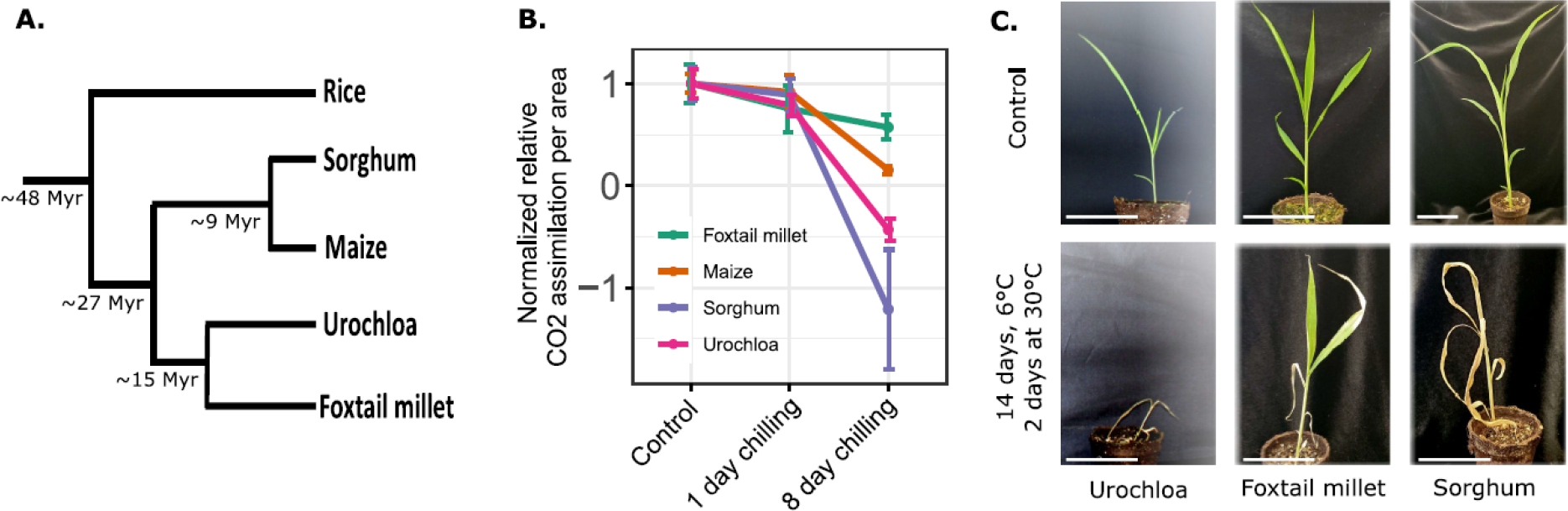
Foxtail millet is a chilling-tolerant representative of the panicoid grass clade. **(A)** Evolutionary relationships of the four species evaluated with rice as an outgroup. Numbers indicate divergence time as reported in Zhang et al. and Pessoa-Filho et al. (23, 58) **(B)** Normalized relative CO_2_ assimilation rates for panicoid grass species with differing degrees of sensitivity or tolerance to chilling stress. CO_2_ assimilation was measured after treatment at 6°C for indicated times below (1 day or 8 days) followed by an overnight recovery of approximately 10 hours at 30°C. Leaf area was measured immediately after assimilation. Individual data points are jittered on the x-axis to avoid overlap. Lines indicate mean values for each species across three replicates and whiskers represent standard error of the mean. **(C)** Phenotypic response of foxtail millet, Urochloa, and sorghum to 6°C chilling stress for 14 days, followed by 2 days of recovery at 30°C. Scale bars, 6cm.

### Foxtail millet membranes have distinct responses to chilling stress

Many cellular membrane systems are damaged in response to chilling (2, 13), and changes in membrane lipid compositions are required to achieve chilling tolerance (7, 30). We profiled membrane lipids from sorghum, Urochloa, and foxtail millet seedlings grown under control and chilling-stress conditions. We hypothesized that patterns unique to foxtail millet and not in both sorghum and Urochloa potentially stem from the difference in chilling tolerance among the species. Likewise, patterns in foxtail millet that are shared by Urochloa but not sorghum are likely to reflect their closer evolutionary relationship. We collected samples for lipid profiling at 10 min, 3 h, 6 h, 12 h, 16 h, and 24 h following onset of chilling-stress. Of the 11 lipids measured (**Table S1**), nine lipids exhibited 24-hour rhythmic accumulation (rhythmic hereafter) in at least one species (**Table S2**) (31). In foxtail millet, all three major membrane lipids, monogalactosyldiacylglycerol (MGDG, LimoRhyde, q-value = 0.07), digalactosyldiacylglycerol (DGDG, LimoRhyde q-value =0.04), and phosphatidylcholines (PC, LimoRhyde q-value =0.03) were found to be rhythmic (Figure 2, Table S2). PC was rhythmic in all three species, while TAG and PG were rhythmic in sorghum and Urochloa. Major lipids such as DGDG and PC were rhythmic in foxtail millet and Urochloa (**Figure 2A, C**), suggesting a strong influence of genetic relatedness on major lipid abundance patterns. However, a foxtail millet-specific increase in MGDG abundance was observed at 24 hr post chilling stress compared to sorghum and Urochloa (p-value = 0.003 and p-value = 0.004, respectively) (**Figure 2A**). Further, we tested the difference in rhythmicity in PC and DGDG between species using CircaCompare analysis (32). The time at which the response variable peaks (phase) is significantly different for PC in all three species (**Table S5**). Mesor, a rhythm adjusted mean, is significantly different in foxtail millet compared to sorghum (p-value = 0.001) and Urochloa (p-value =0.006). These results show that rhythmic lipids across species differ in their peak and average values suggesting species-specific control of rhythmicity in lipid content and composition..

**Figure 2.**
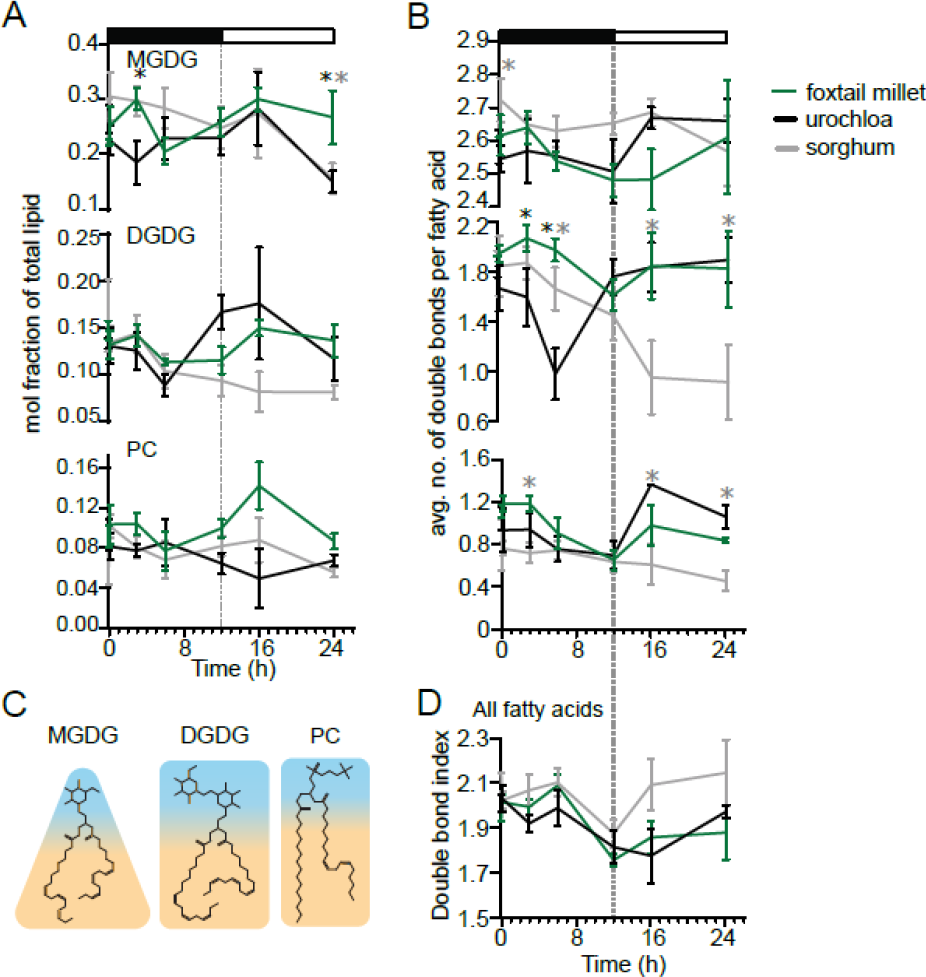
Lipid responses to chilling include effects related to genetic distance and chilling tolerance. The relative abundance of specific lipid species exhibits multiple sequential changes in the first 24 hours of exposure to chilling stress. In all panels, the x-axis indicates the time in hours (h). **(A)** Mole percent abundance of lipids relative to all fatty acid-containing lipids for the following lipid classes: monogalactosyldiacylglycerol (MGDG), digalactosyldiacylglcyerol (DGDG), and phosphatidylcholine (PC) in foxtail millet, Urochloa, and sorghum. **(B)** Unsaturation index, calculated as the average number of double bonds per fatty acid for all fatty acid-containing lipids: MGDG, DGDG, and PC.P-values were determined using Fisher’s least significant difference (LSD). ‘*’ denotes p-value < 0.05, ‘**’ p-value < 0.01, and ‘***’ p-value < 0.001. **(C)** Structural models of major lipids MGDG, DGDG, and PC, where blue indicates the hydrophilic head group and orange indicates the hydrophobic tail group. **(D)** Total unsaturation index, calculated as the average number of double bonds per fatty acid for all fatty acid-containing lipids.

Low temperature induced increases in fatty acid polyunsaturation of membrane lipids are associated with greater membrane fluidity and increased chilling tolerance (10, 33). We detected significant differences in DGDG unsaturation levels in foxtail millet compared to Urochloa following 3 h of chilling stress and relative to Urochloa and sorghum at 6 h of chilling stress, indicating a foxtail millet-specific early stress response (**Figure 2B, Table S4**). We observed similar species-specific differences in lipid abundance and unsaturation levels for minor lipids such as phosphatidylethanolamine (PE), phosphatidylinositol (PI), phosphatidylglycerol (PG), phosphatidylserine (PS), and sulfoquinovosyldiacylglycerol (SQDG) (**Figure S1**). The total lipid unsaturation index remained high for sorghum throughout the time course, while foxtail millet and Urochloa were characterized by lower unsaturation near the end of the time-course (**Figure 2C, Table S4**). Thus, the overall lipid unsaturation in these species cannot explain the increased low-temperature tolerance of foxtail millet.

### Transcriptional changes of lipid metabolism genes are associated with lipid abundance change

In previous work, we have showed that lipid pathway genes were differentially regulated in temperate-adapted *T. dactyloides* compared to maize and sorghum in response to chilling stress and were enriched among genes experiencing rapid rates of protein sequence evolution in *T. dactyloides* (26). To examine whether transcriptional changes in lipid metabolism genes match the observed patterns of lipid changes between chilling-tolerant foxtail millet and chilling-sensitive sorghum, we collected samples from sorghum and foxtail millet for transcriptome sequencing (RNA-seq) at 30 min, 1 h, 3 h, 6 h, 16 h, and 24 h after the onset of chilling stress, as well as from paired control samples not exposed to chilling stress, collected at the same time points. We employed a conventional correlation co-expression clustering method and a linear mixed model (LMM) based method to understand the differences and commonalities in how sorghum and foxtail millet respond to chilling stress at the transcriptional level (see Methods).

We used a set of 16,796 syntenic orthologous gene pairs conserved between sorghum and foxtail millet (25). Of these, 9,778 gene pairs passed an expression data quality filter of standard deviation < 0.4 and r-square > 0.1 (**Table S5**, see Methods). Of this filtered set, 2,233 gene pairs (**Table S6**) exhibited a significant species * treatment interaction effect (multiple testing corrected false discovery rate [FDR] < 0.001 (34), indicating differences in the chilling stress induced transcriptional response of orthologous genes between the two species. In parallel, we applied conventional correlation clustering analysis to identify co-expressed syntenic orthologous gene pairs in sorghum and foxtail millet. We used the ratio of expression values between treatment and control conditions for clustering analysis. Using a permutation test, we defined 16 clusters (see methods; (**Figure S2; Table S7**), and identified 2,245 syntenic orthologous genes in different clusters as being co-expressed orthologs. We classified the remaining 7,533 syntenic orthologs as non-co-expressed orthologs and referred to them as correlation cluster - differentially regulated orthologs (CC-DROs, Table S8). Clusters 2, 4, 6, and 14 had more sorghum genes, while clusters 1, 3, 5, 7, 8, 9, 10, 11, and 13 have higher proportions of foxtail millet genes. Clusters 12, 15, and 16 had similar number of genes from sorghum and foxtail millet (**Table S7**). Clusters 1, 3, 6, and 7 contained genes up-regulated at 6 hr into stress, indicating a possible role in early chilling-stress response. We illustrate the divergence in transcriptional responses to chilling between syntenic gene pairs in a Circos plot, in which lines that cross over between groupings in the center of the chart represent genes that are syntenic orthologs and have distinct patterns of gene expression between foxtail millet and sorghum (**Figure S2**).

We then identified high-confidence differentially regulated orthologs (HC-DROs) by taking the overlap of CC-DROs identified by the clustering method and the DROs identified with LMM (**Table S9**). We determined that 1,708 syntenic orthologous gene pairs overlap in the two sets, which we further used for gene ontology term enrichment (GO) analysis. GO analysis of these 1,708 HC-DRO pairs revealed enrichment for two GO categories: ‘stress response’ and ‘macromolecule metabolic process’. In validation of our focus on lipids, we observed an enrichment for the GO metabolic process category, ‘lipid metabolic process’ (GO:0006629, p-value=0.003, **Table S10**).

We then defined a set of *a priori* candidates from the most likely set of Arabidopsis (*Arabidopsis thaliana)* lipid genes corresponding to fatty acid and glycerolipid metabolism from the AraLipid database (http://aralip.plantbiology.msu.edu/pathways/pathways), and a corresponding set of 356 sorghum-foxtail millet gene pairs homologous to these Arabidopsis genes with syntenic orthologs in both sorghum and foxtail millet. (**Table S11**). The overall gene expression patterns of these 356 gene pairs revealed that lipid-related genes are mostly up-regulated under chilling treatment in chilling-tolerant foxtail millet, but not in sorghum. Of the 356 lipid-related gene pairs, 34 showed differential responses to chilling stress between sorghum and foxtail millet, with pronounced up-regulation of lipid-related gene expression in foxtail millet exposed to chilling stress (**Figure 3, Table S12**). One example of such a differentially regulated ortholog in sorghum and foxtail millet is provided by *3-KETOACYL-COA SYNTHASE 1 (KCS1)*, encoding an enzyme in the fatty acid elongation pathway for wax biosynthesis and involved in chilling tolerance in Arabidopsis (35). The sorghum ortholog of *KCS1*, Sobic.001G438100, was down-regulated throughout the chilling-stress time course. However, the *KCS1* ortholog in the chilling-tolerant foxtail millet, Seita.9G470700, was upregulated at later time points (**Figure 3**), suggesting that the differential regulation of *KCS1* ortholog expression between sorghum and foxtail millet may be leading to the differences in chilling tolerance between the two species.

**Figure 3.**
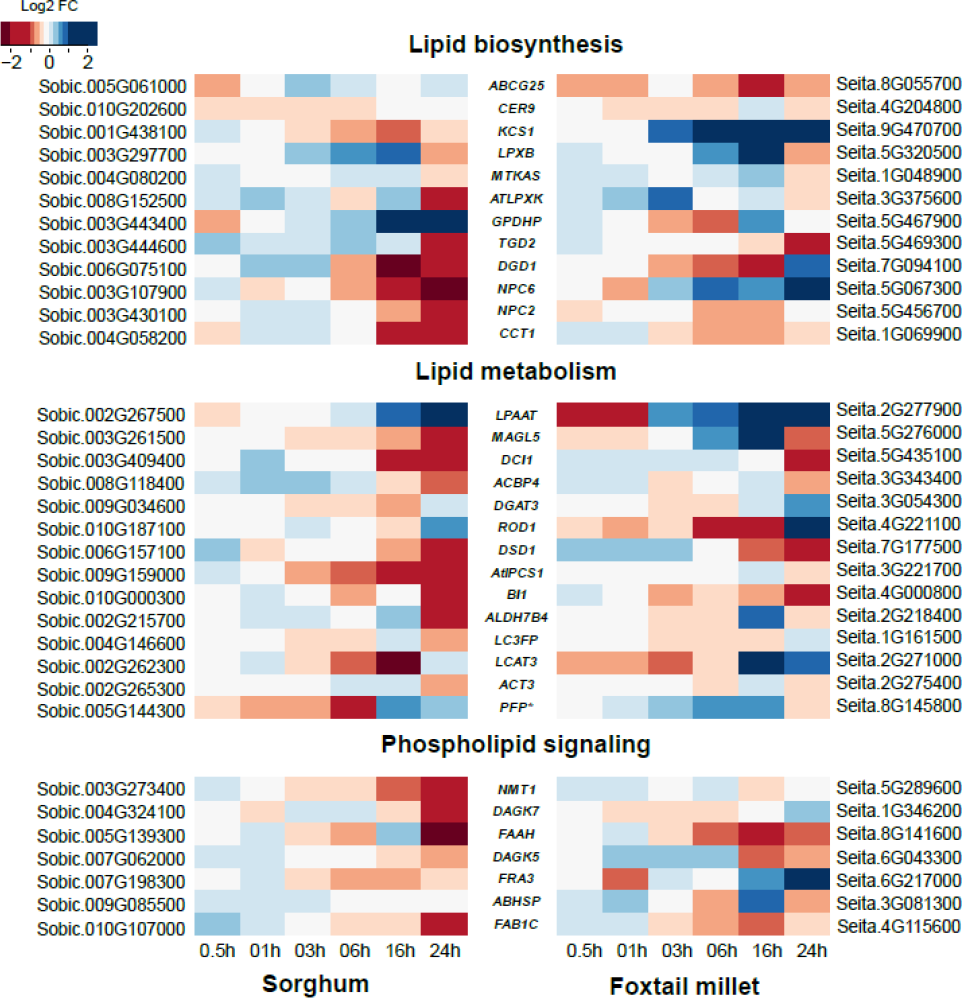
Syntenic orthologs in sorghum and foxtail millet show differential regulation during chilling stress. Heatmap representation of log2 fold change values for chilling stressed samples compared to control in sorghum and foxtail millet at different time points. Lipid related gene pairs that overlapped with high-confidence differentially regulated orthologs were considered and classified into lipid biosynthesis, lipid metabolism, and phospholipid signaling. Genes names in the center are suggestive only and derived from best-hit Arabidopsis genes.

### Gene expression correlation with lipid buildup and breakdown

We asked whether changes in the expression of genes in lipid pathways in foxtail millet and sorghum were consistent with changes in lipid abundance and saturation. To this end, we combined time-course lipid and gene expression profiles to understand how differential gene expression in these two species affects lipid abundance and saturation under chilling stress, using only shared time-points between the two datasets. A diagram of the glycerolipid biosynthesis pathway is shown in **Figure 4A**. Looking at the sorghum ortholog of Arabidopsis *DIGALACTOSYL DIACYLGLYCEROL DEFICIENT 1 (DGD1)*, Sobic.006G075100, its expression profile had significant and positive correlation with DGDG accumulation during chilling (Pearsons correlation coefficient (PCC r = 0.85, p-value = 0.03). However, the expression of the *DGD1* ortholog in foxtail millet was not correlated with DGDG accumulation (**Figure 4B**). Similarly, the expression of *NON-SPECIFIC PHOSPHOLIPASE C1 (NPC1)* orthologs in sorghum and foxtail millet was positively correlated with PE accumulation during chilling (PCC r = 0.79, p-value = 0.06; PCC r =0.84, p-value = 0.03, respectively). However, the expression of *NPC2, NPC6*, and *PHOSPHOLIPID N METHYLTRANSFERASE* (*PLMT)* was also positively correlated with PE accumulation in sorghum but not in foxtail millet (**Figure 4C**). Notably, we detected a significant correlation between gene expression and lipid contents for lipids with significant species-specific changes in lipid abundance, such as MGDG and DGDG, as shown in Figure 2 and Figure 4B.

**Figure 4.**
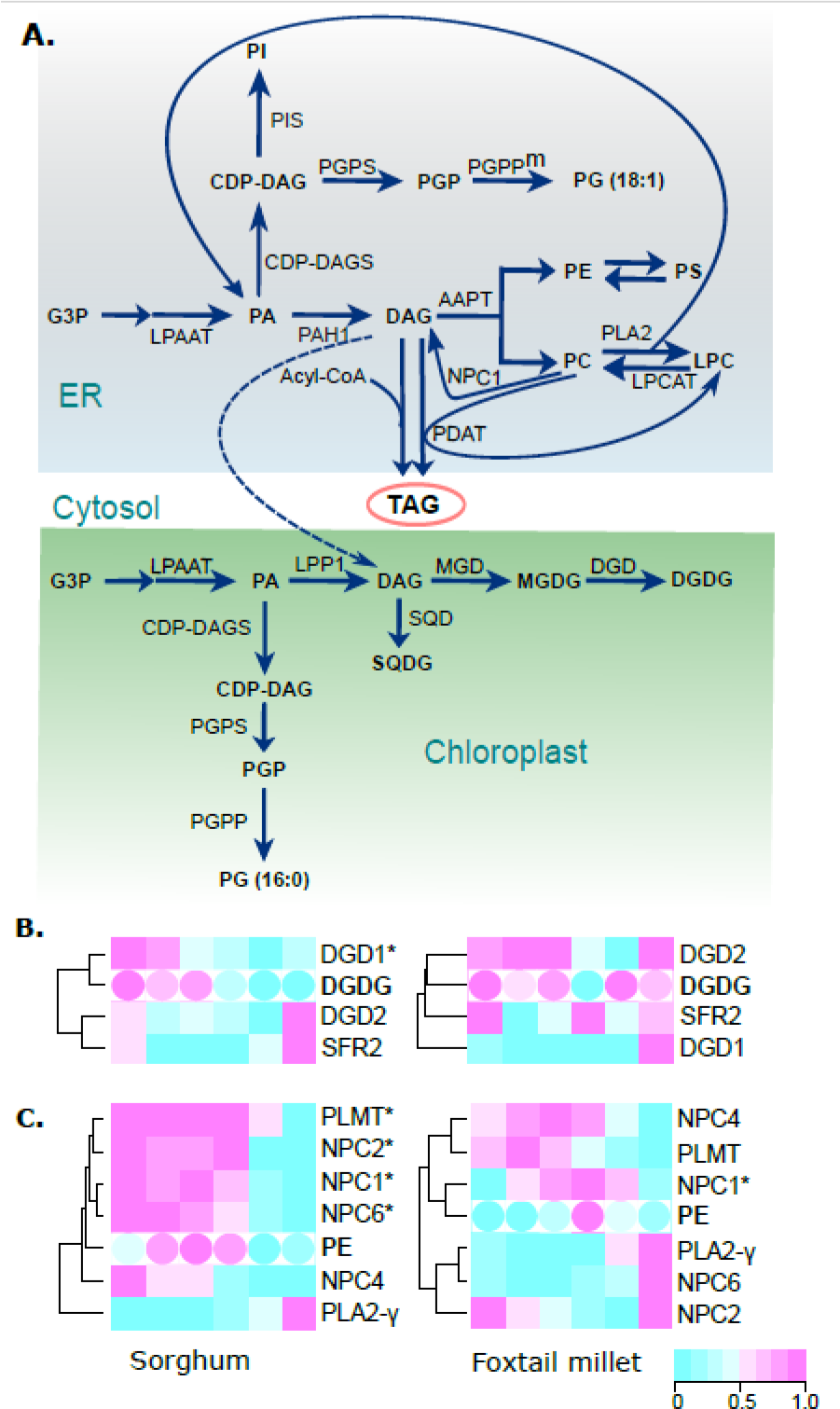
Correlation of lipid and transcript abundances in the glycerolipid biosynthesis pathway allows identification of candidate genes. **(A)** Diagram of the glycerolipid biosynthesis pathway with lipid species shown in bold font and the enzymes responsible for each step denoted next to the corresponding arrows. **(B, C)** Heatmaps showing transcript abundance for candidate genes in lipid metabolism and abundance of lipid species (in bold font) at different time points for DGDG **(B)** and phosphatidylethanolamine (PE) **(C)**. Significant correlations measured by pearson correlation, between lipid changes and transcript abundance in either sorghum or foxtail millet are indicated by ‘*’ after the gene name.

The accumulation of triacylglycerols (TAGs) in plants arises from multiple sources (36, 37) and TAGs are important for low-temperature tolerance (38–40). Correlating gene expression patterns with TAG abundance was expected to shed light on the potential source of TAG during the chilling response. The expression levels of the foxtail millet ortholog to Arabidopsis *LIPID PHOSPHATE PHOSPHATASE 2 (LPP2)*, Seita.4G217800, showed a significant and positive correlation with lipid abundance in TAG accumulation during chilling stress response in foxtail millet (PCC r = 0.86, p-value = 0.003). By contrast, the expression levels of the sorghum ortholog to LPP2, Sobic.010G190300, showed no significant correlation with TAG accumulation (**Table S13**). This finding suggests that, at least in foxtail millet, phospholipids are the primary source of chilling-stress-induced TAG accumulation. A list of specific orthologs in foxtail millet and sorghum whose expression levels were significantly correlated with the buildup and breakdown of each lipid species is provided in **Tables S13 and S14**.

### Conservation of chilling-induced changes in lipid composition and rhythmicity in Arabidopsis

The amplitude of changes in lipid level observed in panicoid grasses in the first 24 hours of chilling was beyond our expectations. To test if this amplitude is evolutionarily conserved beyond the grasses, we quantified representative lipids from Arabidopsis seedlings across a time-course with paired control samples and chilling-stress samples collected immediately before chilling stress (0 h), and after 2 h, 6 h, 10 h, 14 h, 18 h, 22 h, and 26 h of stress. DGDG levels remained constant during normal conditions or chilling stress, whereas MGDG levels were slightly higher upon chilling stress compared to control conditions, reaching statistical significance at 22 h and 26 h into stress (**Figure 5A**). This increase in MGDG levels at the late time points was similar to the significant increase in MGDG after 24 h of exposure to stress in chilling-tolerant foxtail millet compared to chilling-susceptible sorghum and Urochloa (**Figure 2**). PC levels increased and remained higher across the entire time course (**Figure 5A, Table S15**). However, PC unsaturation under chilling stress conditions was only distinguishable from control samples at a few time points (**Figure 5B, C**). We detected significant rhythmicity in MGDG levels in control conditions (Table S16, rhythmic p-value = 0.005) using the ‘circa_single’ method in CircaCompare analysis (32). However, the amplitudes of major lipids - MGDG, DGDG, and PC were much lower in Arabidopsis compared to the three grasses. These results suggest the conservation of chilling tolerance-induced changes in lipid content and composition and rhythmic patterns of lipids across grasses and Arabidopsis despite 150 million years of divergence between monocots and eudicots (41).

**Figure 5.**
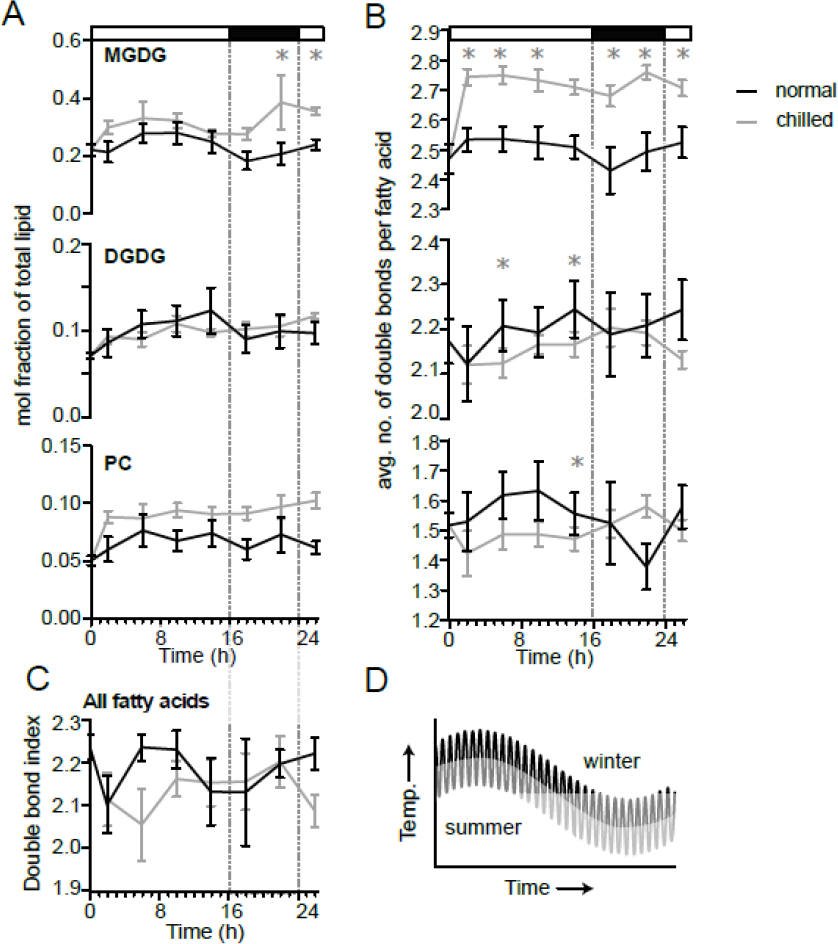
Time-series profiles of Arabidopsis lipid abundance and unsaturation during chilling or control conditions. **(A)** Mole percent abundance of lipids relative to all fatty acid-containing lipids for the following lipid classes: monogalactosyldiacylglycerol (MGDG), digalactosyldiacylglcyerol (DGDG), and phosphatidylcholine (PC) in Arabidopiss during chilling or control conditions. **(B)** Unsaturation index, calculated as the average number of double bonds per fatty acid for MGDG, DGDG, and PC. **(C)** Unsaturation index, calculated as the average number of double bonds per fatty acid for all fatty acid-containing lipids. Significant p-values determined using Fisher’s least significant differences. *’ denotes p-value < 0.05, ‘**’ p-value < 0.01, and ‘***’ p-value < 0.001 **(D)** Diagram showing daily and seasonal fluctuation of temperature during the plants’ growth cycle.

#### Lipid-related genes exhibit expression rhythmicity

To determine which sorghum and foxtail millet lipid related genes exhibit 24-hour rhythms, we examined the patterns of 356 sorghum-foxtail millet lipid related gene pairs in a previously published 72 h RNA-seq time-sourse (42). We identified 224 sorghum and 189 foxtail millet genes in this set as being rhythmically expressed. Of these, 145 pairs were rhythmic in both sorghum and foxtail millet. We then used the *LimoRhyde* package (31) to identify those genes with rhythmic expression under control conditions in our data sets. This analysis indicated that 131 sorghum and 204 foxtail millet lipid related genes had rhythmic expression patterns under control conditions (**Table S17**). Further, we employed LimoRhyde to test for differences in rhythmic expression, or differential rhythmicity (DR), for each gene between the control and chilling treatments in sorghum and foxtail millet. We identified 142 foxtail millet lipid related genes and 101 sorghum lipid-related genes displaying differential rhythmicity between the control and chilling treatments (**Table S17**). Among the 58 lipid related gene pairs that showed DR between control and chilling stress in both sorghum and foxtail millet, 36 showed rhythmic expression under control conditions in both species (**Figure 6A**). These lipid related gene pairs are rhythmic genes that change their rhythmicity patterns under chilling treatment and likely represent shared targets in sorghum and foxtail millet for chilling stress induced alterations in expression.

**Figure 6.**
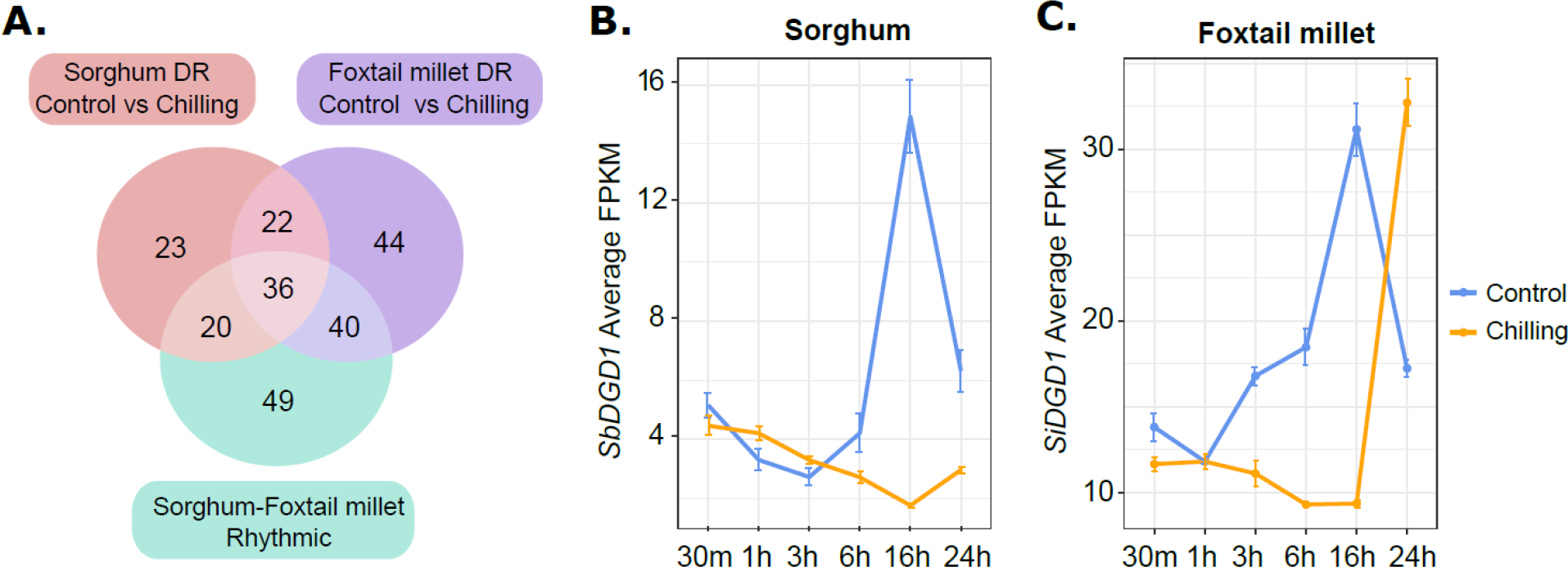
Comparison of rhythmicity and differential rhythmicity in sorghum and foxtail millet lipid related genes. **(A)** Venn diagram showing the extent of overlap between rhythmic genes in sorghum and/or foxtail millet identified by *LimoRhyde* analysis under control conditions (244 genes in the green circle) and sorghum differentially rhythmic (DR) under control conditions compared to chilling treatment (101 genes in the pink circle) and foxtail millet DR under control conditions compared to chilling treatment (142 genes in the purple circle). The union of these three data sets is 36 genes representing rhythmic genes under control conditions that change their rhythmicity in response to chilling temperature in both foxtail millet and sorghum. **(B**,**C)** An example of a rhythmic gene *DGD1* showing a similar diurnal expression pattern under control conditions and changed rhythmicity during chilling stress response in sorghum **(B)** and foxtail millet **(C)**. *DGD1* expression is shown as average FPKM values from three biological replicates and the standard error of mean.

An example of such a lipid-related gene whose rhythmic expression under control conditions is altered during chilling stress in both sorghum and foxtail millet was *DGD1* (**Figure 6B, C**). Sorghum and foxtail millet *DGD1* showed similar rhythmic patterns of expression under control conditions. However, their rhythm and magnitude of expression change significantly during chilling stress in both species. Of note, *DGD1* was identified as a differentially regulated ortholog in **Figure 3**. In addition, *SbDGD1* expression during chilling stress was positively correlated with DGDG abundance in sorghum. The foxtail millet *DGD1* ortholog did not show such a correlation, suggesting species-specific changes in their response to chilling stress. These results indicate that differences in the diurnal regulation of lipid-related genes in sorghum and foxtail millet may lead to differential responses to chilling stress.

## DISCUSSION

Panicoid grasses represent an interesting clade with repeated gain or loss of chilling tolerance, reflecting parallel adaptation strategies in different lineages within the clade. Using representative chilling-susceptible sorghum and chilling-tolerant foxtail millet allowed us to identify changes to transcript and lipid levels that are likely to be functionally linked to variation in chilling tolerance between the related species. We included Urochloa, a chilling-sensitive panicoid grass that is more closely related to foxtail millet, as a control for the large evolutionary divergence between foxtail millet and sorghum.

Here, we assembled time course datasets for transcript levels and lipid metabolic profiling in three panicoid grasses with different genetic relatedness and tolerance to chilling stress to understand whether and how changes in composition of membrane lipids and corresponding changes in gene expression contribute to chilling tolerance in foxtail millet. Most changes in lipid content and composition were consistent across the three species, likely representing shared responses to chilling stress due to their genetic relatedness. Lipid unsaturation appears to be the first response to a need for increased membrane fluidity as the temperature decreases (i.e., lipid unsaturation at 10 min time point is high in all species, **Figure 2D, Table S1, S15**). Still, the degree of lipid unsaturation did not increase throughout the time course, again across all species. Urochloa appears to behave more like foxtail millet than it does like chilling-susceptible sorghum, likely an effect of its short evolutionary distance compared to foxtail millet. By comparing sorghum to Urochloa, we were able to tease out a small subset of lipid metabolic changes that are unique to foxtail millet, the most chilling tolerant panicoid grass tested in this study (**Figure 2**). These results also indicate that lipid unsaturation is unlikely to be the source of chilling tolerance in foxtail millet, as it is similarly adjusted in all three species during the first 24 hours of chilling (**Figure 2B, S2**).

There is little consensus in reports of changes in lipid content and composition in response to cold stress across land plants (14). These discrepancies in lipid-related changes may reflect inherent genetic and physiological differences in how individual species respond to chilling stress; alternatively, they may stem from varying experimental designs and variation due to sampling time. Evidence is fast emerging for the role of circadian clock regulation in coordinating dynamic plant responses to daily and seasonal environmental fluctuations (43, 44). However, daily rhythms in lipid metabolism had not previously been reported for important clades of crops like panicoid grasses under chilling conditions. Notably, rhythmic changes in lipid composition and gene expression during chilling stress are not similar across species, suggesting that a general strategy is not to stop or slow down the circadian clock during stress, rather plants may have developed species-specific strategies to overcome these challenges. Our time-series dataset of changes affecting lipid during chilling stress in three grasses allowed us to uncover rhythmic patterns of lipid abundance and unsaturation. Moreover, our lipid dataset reveals chilling-tolerance-related changes in lipid abundance and unsaturation at specific time points, potentially explaining part of the difficulty in extracting conserved patterns for lipids across species in previous reports involving one or a few time points. We also detected rhythmicity in expression of lipid metabolic genes in both sorghum and foxtail millet. The rhythmic nature of changes in lipids and specific changes during chilling stress were similar between grasses and Arabidopsis, in contrast to other published studies that show differences. These findings highlight the importance of time-series datasets to account for diurnal cycles in uncovering conserved features of chilling-stress responses across large phylogenetic distances. While the changes in lipid content and composition in Arabidopsis are consistent with the panicoid grasses, the amplitude of changes in lipid abundance in grasses is about twice that of Arabidopsis (45, 46). This lower amplitude of major lipids could also explain our inability to detect consistent rhythmicity in all major lipids in Arabidopsis. More detailed analysis of lipid measurements across genotypes and species under chilling stress conditions is needed to understand the variation in amplitudes of changes across species more thoroughly.

Combined lipidomic and transcriptomic analysis has been used to unravel transcriptional regulation of lipid metabolism during chilling-stress responses in maize (47). Our results show species-specific differences in transcriptional correlation with lipid metabolic changes, suggesting complex regulation of metabolic perturbations involved in plants’ response to environmental challenges. We propose that this approach with chilling-susceptible and chilling-tolerant species can empower the identification of specific genes whose transcript levels are correlated with changes in lipid metabolites, in response to chilling stress. Although, our experimental design may miss non-syntenic genes that may have acquired novel chilling-induced changes in their expression. We identified species-specific differences in the extent of correlation between lipid related gene expression and changes in lipid abundance changes, potentially informing the flux of fatty acids. For example, we discovered that the expression of the foxtail millet *LPP2* is tightly correlated with TAG abundance. This correlation indicates that phospholipids are the primary source of TAG in foxtail millet during chilling response. By contrast, the expression of the sorghum ortholog of *LPP2* was not correlated with TAG abundance, suggesting species-specific differences in TAG accumulation or that the primary source of TAG differs between these two species during chilling stress.

*NPC1* expression and PE levels are another example of an unexpected lipase influencing lipid levels. *NPC1* expression was correlated with PE accumulation in foxtail millet and sorghum. *NPC1* produces DAG either through the hydrolysis of PC or MGDG and DGDG, which can in turn be converted to PE (48, 49). The role of *NPC1* in response to heat stress is known (49). Here, we propose a role for *NPC1* in PE accumulation and chilling tolerance in panicoid grasses.

Overall, we show that despite the conservation of many transcriptional and metabolic responses to chilling stress across species, the unique combination of species employed in our study allowed us to identify a smaller set of genes more likely to be functionally linked to variation in chilling tolerance than merely due to genetic relatedness. This study provides a framework to probe potential genes whose function in changes to lipid content and composition may not be previously known. For the first time, we also report diurnal rhythmicity in lipid abundance, saturation, and expression of lipid-related genes in these panicoid grasses during chilling stress.

## Methods

### Plant growth and chilling treatment

Seeds for the reference genotypes for sorghum (*Sorghum bicolor, BTx623*), maize (*Zea mays, B73*), Urochloa (*Urochloa fusca, LBJWC-52*) and foxtail millet (*Setaria italica,Yugu1*) were grown in a Percival growth chamber (E-41L2) with 60% relative humidity, with a 12 h light/12 h dark photoperiod and a target temperature of 29°C during the day and 23°C at night. Chilling stress was applied to 12-day-old seedlings, when all species were developmentally equivalent based on number of leaves. Immediately at the end of the light period, seedlings were moved to a second growth chamber with equivalent growth settings except for a target temperature of 6°C. Each sample represents a pool of above-ground tissue from at least three seedlings. Samples were harvested from the control conditions and chilling stress treated plants at the designated time points after the onset of chilling stress.

For lipid analysis, samples were harvested at 0 min (immediately before reducing the chamber temperature to 6°C), 10 min, 3 h, 6 h, 12 h, and 24 h. The 10 minute sample was taken 10 after the chamber air temperature reached 6°C, approximately 20 minutes past time 0. Whole shoot tissue excluding the coleoptile, was collected at the soil level. The tissue was quickly and gently submerged in 1 mL of ice-cold extraction solvent (2:1:0.1 v/v/v methanol:chloroform:formic acid) in a 2 ml tube and shaken on a bead beater at 4000 inversions per minute in 30-second intervals with intervening ice incubations until the tissue was thoroughly disrupted. Lipid extraction continued following a modified Bligh and Dyer protocol (50). Following extraction, lipids were concentrated and stored at -80°C under nitrogen. Lipids were separated as described in Wang and Benning (51) with the following modifications. A 10% lipid spot was loaded in the corner of each thin layer chromatography (TLC) plate that did not see solvent which served as a control for total fatty acids, and was used to make internal comparisons.

A two-dimensional TLC plate was used for separation. In the first dimension, a mixture of chloroform: methanol: ammonium hydroxide, (130:50:10, v/v/v) was used as solvent and in the second dimension, chloroform: methanol: acetic acid: water (85:12.5:12.5:4, v/v/v/v) was used as a solvent. A separate one dimensional thin-layer chromatogram was used to separate non-polar triacylglcyerol, with petroleum ether:diethyl ether:acetic acid (80:20:1, v/v/v) as solvent. Lipids were identified by retention time compared to standards purchased from Avanti Polar Lipids. Remaining analysis was precisely done as described in Barnes et al 2016 (12)

The statistical analysis of lipid data involved several steps. Outliers were assessed at two levels. Firstly, for fatty acid abundance, a robust regression of outlier removal (ROUT) analysis was performed at a 10% threshold using GraphPad v9.5.0 to eliminate any misidentified peaks or anomalies. Any outliers detected at this step were removed from further analysis. Second, the relative mole percentages of each lipid were calculated and normalized to the total fatty acids present. The resulting mole percentages were then screened for outliers using one interquartile distance from the median for each lipid class for each genotype at each temperature. Asterisks denote statistical significance (p ≤ 0.05), determined by fitting a mixed model, with Tukey’s correction for multiple tests.

The double bond index (DBI) was calculated using the formula: ((X:1)×1+(X:2)×2+(X:3)×3)/100, where X represents the relative mole % of 16:n and 18:n fatty acids, and n denotes one, two, or three double bonds. Multiple comparisons were adjusted using Tukey’s multiple comparisons test when comparing between genotypes. Each analysis was performed on at least two growth trials and three biological replicates.

### Measurement of CO_2_ assimilation rates

Seedlings were grown and stress treated as above, with the modification that small plastic caps were placed over sorghum, foxtail millet, and Urochloa seedlings to prevent them from becoming too tall to fit into the LI-COR measurement chamber. After 0, 1, or 8 days of chilling treatment, seedlings were allowed to recover in the greenhouse overnight under control conditions and CO_2_ assimilation rates were measured the next morning using the LI-6400 portable photosystem unit under the following conditions: PAR 200 μmol mol−1, CO_2_ at 400 μmol mol−1 with flow at 400 μmol mol−1 and humidity at greenhouse conditions. Whole seedlings readings were measured for sorghum, foxtail millet, and Urochloa after covering the pots with clay and using the LI-COR’s “Arabidopsis chamber.” Readings for maize were measured using the leaf clamp attachment which was always placed on the second leaf at a position 3 cm above the ligule. Leaf area was measured using the LI-3100C area meter.

### RNA isolation and RNA-seq analysis

Total RNA was isolated from paired samples collected at 30 min, 1 h, 3 h, 6 h, 16 h, and 24 h after the onset of chilling. Library construction was performed following the protocol described by Zhang et al. (25). Sequencing was conducted at the Illumina Sequencing Genomics Resources Core Facility at Weill Cornell Medical College. Raw sequencing data are available through the NCBI (http://www.ncbi.nlm.nih.gov/bioproject) under accession number SRA: SRP090583 and BioProject: PRJNA344653. Summary statistics for all the libraries are provided in Table S1. Adapters were removed from the raw sequence reads using *cutadapt* v*1*.*6*. RNA-seq reads were mapped to genome assemblies downloaded from Phytozome (v13): v3.1 (sorghum) and v2.2 (foxtail millet). RNA-seq reads from each species were aligned using GSNAP (52) and Fragments Per Kilobase of transcript per Million mapped reads (FPKM) values were obtained using cufflinks v2.2.1 (53).

### Syntenic orthologs in sorghum and foxtail millet

A final set of 9778 syntenic orthologous gene pairs between sorghum and foxtail millet was calculated from the previously published list of syntenic orthologs (54) after filtering for standard deviation < 0.4 and r-square > 0.1 of the FPKM values (**Table S5**). Clustering was performed using ‘correlation’ from R packages ‘fpc’ (55, 56). To observe treatment effects, the ratio between treatment FPKM and control FPKM was determined for the time course. A permutation test was performed by keeping the sorghum gene constant and randomly assigning a different foxtail millet gene 100 times to determine the appropriate r2, standard deviation, and number of clusters. Syntenic orthologs found within the same clusters were considered co-expressed (CEO), while syntenic orthologs found in different clusters were considered as differentially expressed orthologs (DEO)

### Identification of differentially regulated orthologs

The FPKM values were measured from three biological replicates each for sorghum and foxtail millet under control and cold treatment at six time points (30 min, 1 h, 3 h, 6 h, 16 h, and 24 h). Similar to the cluster analysis, the treatment over control (T/C) FPKM ratios at each time point for sorghum and foxtail millet were calculated and treated as a response. A linear mixed model (LMM) was used to model the T/C ratios as a relationship between the species identity and sampling time under chilling stress on the same set of syntenic orthologous gene pairs used in the cluster analysis. Let yijkl denote the T/C ratio of the ith gene from the kth species and the lth biological replication at the jth time point, where j = 1-6 to represent the six time points, k = 1, or 2 to represent the two species: sorghum and foxtail millet, and l = 1, 2, or 3 to represent the three biological replicates. There were a total of six biological replicates in the study, three from sorghum and three from foxtail millet. We modeled the biological replication as a random effect. For the ith gene, conditioned on this random replication effect, the response yijkl is normally distributed with mean μijkl and variance σ 2 i. The expected T/C ratio μijkl was linearly related to the species, time and their interactions as μijkl = vi + αij + βik + γijk + ηikl for ηikl ∼ N(0, θ2 i), (1) where vi is the intercept; αij and βik stand for the main effect of time and species for the ith gene respectively; γijk is the interaction between time and species, denoting different patterns of expression between the two species over time; and ηikl is the random effect for the biological replicates, which is assumed to be normally distributed with mean 0 and variance θ 2 i. Note that the interaction effect γijk in the model (1) stands for the difference of the T/C ratios over time between sorghum and foxtail millet. The non-zero interaction effect represents different patterns of T/C ratios changing over time between the two species, while the zero γijk indicates a similar trend of the responses of the two species. Those genes with nonzero interaction effect are defined as differential regulated orthologs (DRO) and the ones with zero interaction effect are called comparable regulated orthologs (CRO). In order to identify the DROs, we considered the hypotheses Hi,0 : γijk = 0 for all j, k vs. Hi,a : γijk 6= 0 for some j, k (2) for each gene.

Estimation of γijk and its associated standard error were obtained by the *‘lmer’* function in the R package *lme4*. Wald test statistic was conducted for the hypothesis (2), and the associated p-value for each gene was calculated. Benjamini and Hocheberg multiple test correction was applied to control for false discovery rates (FDR) > 0.001. Those pairs with FDR < 0.001 were considered as DRO, and those with FDR > 0.01 were considered as CROs.

### Lipid genes in sorghum and foxtail millet

A manually curated list of Arabidopsis genes known to be involved in lipid pathways was first created using the Aralip database (http://aralip.plantbiology.msu.edu/pathways/pathways). The sorghum and foxtail millet genes were then matched to the Arabidopsis lipid genes using the published best Arabidopsis hits for the sorghum and foxtail millet genome on Phytozome (v13). Each sorghum and foxtail millet hit was matched with their respective syntenic ortholog in the other species, creating a list of syntenic orthologous pairs of lipid genes in sorghum and foxtail millet (Table S9).

### Gene expression and lipid heatmaps

FPKM values and lipid abundance and unsaturation were normalized by linear transformation such that the minimum value within the time series turned into a zero and maximum values are turned to one. All other values get transformed into decimals between 0 and 1. Heatmaps were generated using heatmap2 function in R.

### Identification of rhythmicity in lipid abundance and expression of lipid related genes

Rhythms in lipid abundance were identified with the ‘circa_single’ method in CircaCompare (package version 0.1.1) in R (version 4.3.0) with default settings (32). Differences in lipid abundance waveforms were detected with the ‘circacompare’ method in the same package. FPKM values representing expression at 3-hour intervals over 72-hours for the 356 lipid-metabolism-associated genes that were syntenic between sorghum and foxtail millet were derived from previously published transcriptomes of comparably staged third-leaf-stage seedlings from sorghum, foxtail millet, and maize (42). Genes in the 356 metabolism-associated data set exhibiting differential rhythmicity between temperature treatments (i.e., cold treatment vs. no treatment) or genotypes (sorghum vs. foxtail millet) were identified with the R package *LimoRhyde* (31) in Bioconductor (57). LimoRhyde reports Benjamini and Hochberg q-values (34) of the rhythmicity of gene and diifferential rhythmicity for genes shared between the two data sets. Statistical significance for either rhythmicity or differential rhythmicity was set at a q-value of ≤ 0.05. Foxtail millet genes were keyed to their sorghum synteologs for LimoRhyde identification of differential rhythmicity between sorghum and foxtail millet genes.

