## Supplementary material for "Rhythmic lipid and gene expression responses to chilling in panicoid grasses": Figure S

**Figure S1: Minor lipid responses to chilling include effects related to genetic distance and chilling tolerance.**

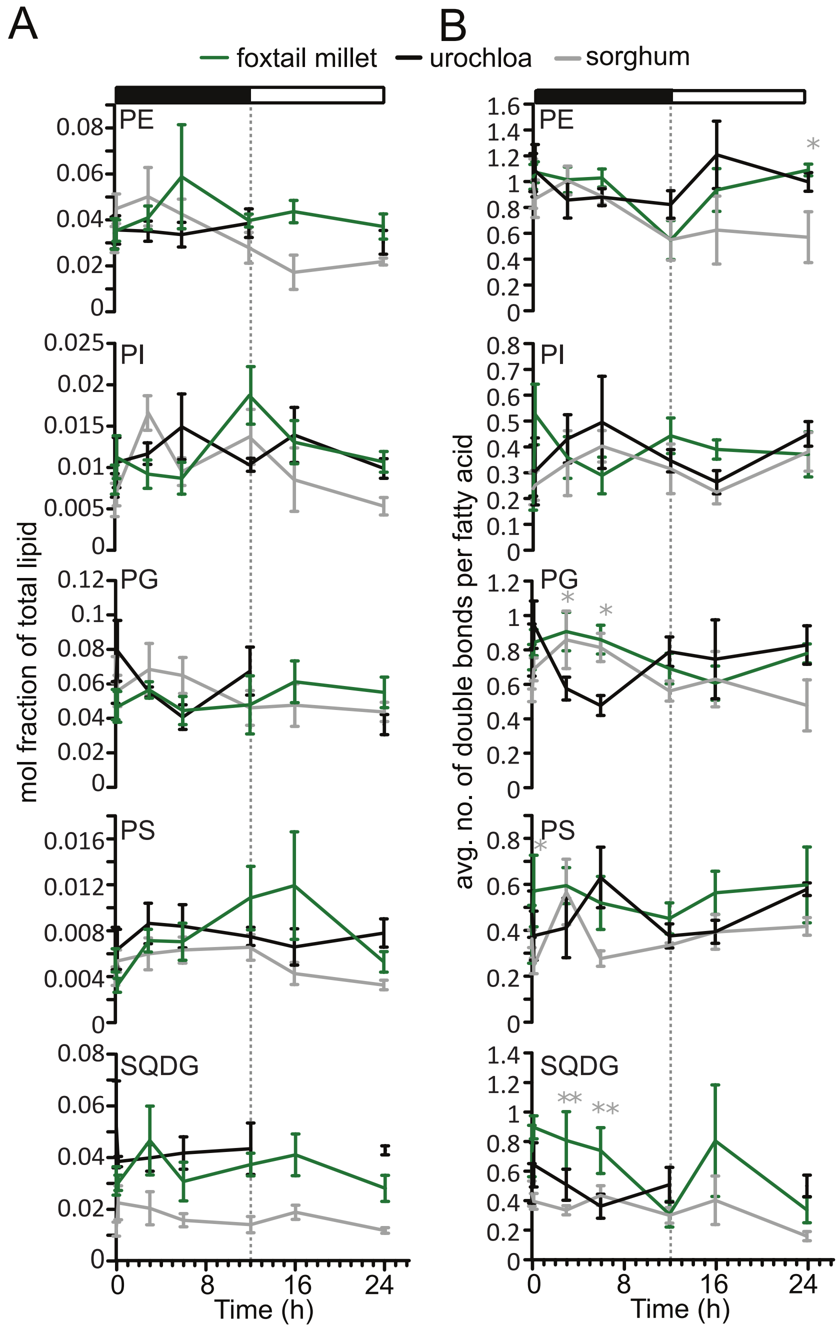

**Figure S2: Circos plot showing differential expression of sorghum - foxtail millet orthologous genes.**
Genes were grouped into 16 co-expression clusters based on patterns of transcriptional abundance. The average pattern of expression across six time points (0.5h, 1h, 3h, 6h, 16h, and 24h) for each cluster is shown in the outermost ring, with the pattern observed in each of three biological replicates drawn separately in black, green, or red. Expression patterns for individual genes within a cluster are indicated in red and blue, with 0.5 hours as the innermost of six layers and 24 hours the outermost of the six layers. The size of the colored bars indicates the number of genes assigned to each cluster in each species, with foxtail millet coming after sorghum in a clockwise progression around the figure. The thickness of connecting lines in the center of the figure indicate the frequency which with sorghum and foxtail millet orthologous gene pairs are assigned to different pairs of co-expression clusters. Lines were only drawn between clusters which shared at least 5% of total assigned gene pairs..

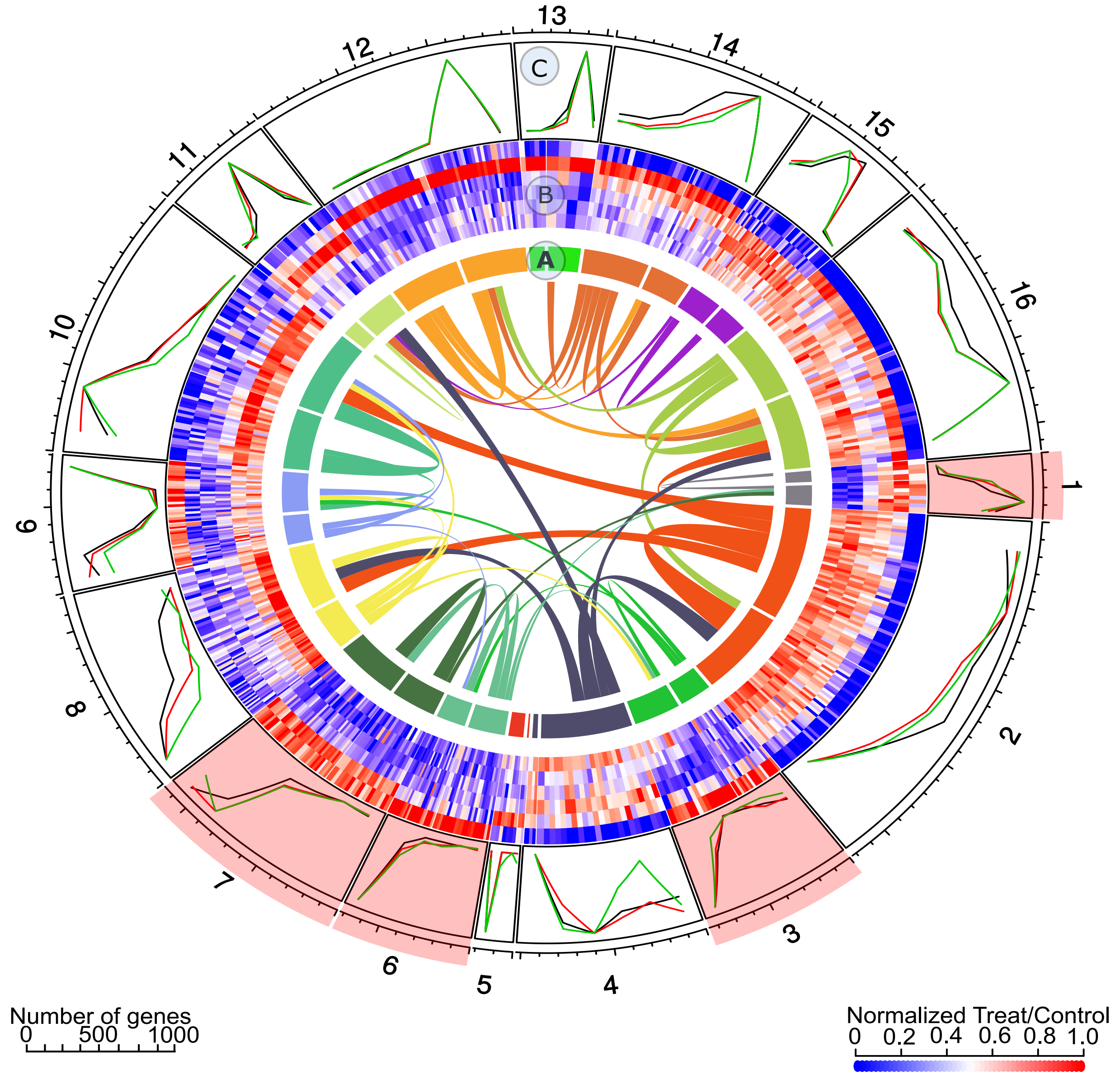
